# Predicting biological pathways of chemical compounds with a profile-inspired aproach

**DOI:** 10.1101/2021.03.02.433511

**Authors:** Javier Lopez-Ibañez, Florencio Pazos, Monica Chagoyen

## Abstract

Assignment of chemical compounds to biological pathways is a crucial step to understand the relationship between the chemical repertory of an organism and its biology. Protein sequence profiles are very successful in capturing the main structural and functional features of a protein family, and can be used to assign new members to it based on matching of their sequences against these profiles. In this work, we extend this idea to chemical compounds, constructing a profile-inspired model for a set of related metabolites (those in the same biological pathway), based on a fragment-based vectorial representation of their chemical structures. We use this representation to predict the biological pathway of a chemical compound with good overall accuracy (AUC 0.74-0.90 depending on the database tested), and analyzed some factors that affect performance. The approach, which is compared with equivalent methods, can in addition detect those molecular fragments characteristic of a pathway. The method is available as a graphical interactive web server http://csbg.cnb.csic.es/iFragMent

## Introduction

Studying the roles of chemical compounds in a cellular context is fundamental for understanding living systems at the molecular level [1]. This can be achieved with experimental and computational approaches. Among the last, of special interest is the analysis of relations between the structure of a chemical compound and its biological role.

Predicting the biological role of a chemical compound, i.e. the pathway(s) it is involved in, from its chemical structure would be valuable in many aspects, ranging from the identification of compounds in metabolomics experiments to the prediction of possible biological roles of drugs. A number of studies aimed to predict the biological pathway of a compound from its 2D structure alone. Most biological pathways contain a reduced number of metabolites, what could be a problem for machine learning approaches. This is one of the reasons why most studies have tried to predict general pathway classes [2–7] (e.g. carbohydrate metabolism). While predicting the general pathway class of a compound would be valuable in some cases, most applications will require the prediction of specific pathways (e.g. glycolysis).

Two previous studies, in addition to predict general pathway classes, have also tried to predict specific pathways. In the context of a wider analysis of human metabolic pathways and their mapping in the chemical space, Macchiarulo *et al.* [8], using a machine learning algorithm (random forests), assigned compounds to 52 human metabolic pathways defined in KEGG [9]. More recently, Hamdalla *et al.* [10], proposed a family of related approaches based on a ranking algorithm and pair-wise substructure matching that were tested in 137 KEGG metabolic pathways. Their best performing approach (implemented as a software package, TrackSM) followed a two-step classification: in the first step, TrackSM predicts the pathway class of a compound, and in a second step it identifies the specific pathway from that class.

In this work we propose a method, iFragMent, for the prediction of compounds involved in specific pathways (not pathway classes), inspired by sequence profile approaches. Sequence profiles have been widely used in the study of DNA and protein sequences, and they are behind most modern methodologies for obtaining information from these biological polymers. Sequence profiles are formal models that capture the main characteristics of a set of related sequences. They are built from a multiple sequence alignment of these related sequences using different approaches, from simple “position specific scoring matrices” (PSSMs) to complex statistical models such as “hidden Markov models” (HMMs) [11]. Once built, these profiles allow assigning new members to the family (hence predicting their function if it is unknown), detect functionally important residues (e.g. conserved positions) or define domains, among other things. These profile-based approaches have been designed taking into account the polymeric nature of DNA and proteins and their underlying evolutionary relationships.

We explore the possibility of using a conceptually (not methodologically) related strategy for studying chemical compounds in the context of biological pathways. We take into account the chemical composition (in the form of chemical fragments) of the whole set of compounds participating in a pathway (e.g. metabolic, regulatory and signalling networks). We calculate the enrichment of chemical fragments in the pathway, and use this information to score new compounds, given the presence of the pathway-enriched fragments in their structure. We evaluate our method in its ability to predict the correct pathway of a chemical compound, using 861 pathways defined in four databases: KEGG [9], Reactome [11], SMPDB [12] and enviPath [13].

Our results show that the method proposed predicts biological pathways for chemical compounds with global AUCs ranging from 0.74 to 0.90 depending of the database considered. We compare our method to previous aproaches [8,10] and to a k-nearest neighbor approach based on pair-wise structural similarities. In addition to the predicted pathway, our approach reports associated p-values and detects the chemical substructures responsible for a compound-pathway assignment. The method is implemented as a web server http://csbg.cnb.csic.es/iFragMent

## Methods

### Datasets

We compiled pathways and their associated compounds from four resources: KEGG (“pathway” section, release 83.0), Reactome (version v61), SMPDB (release 2.75) and enviPath (EAWAG-BBD dataset, version 0.3.1). Compounds were compiled regardless the organism they are eventually assigned to. We selected those pathways with at least ten chemical compounds, excluding the very general pathways (e.g. from KEGG ‘Metabolic pathways’ (map01100); ‘Biosynthesis of secondary metabolites’ (map01110); ‘Microbial metabolism in diverse environments’ (map01120); ‘Biosynthesis of antibiotics’ (map01130) and ‘Degradation of aromatic compounds’ (map01220)).

Secondary structure files of compounds were downloaded from their respective database for KEGG, EnviPath and SMPDB pathways. For Reactome pathways, we retrieved chemical structure files from ChEBI [15] using the cross-references provided in Reactome. Datasets (excepting KEGG, due to license requirements) are available at https://github.com/jlopez-ibanez/iFragMent

### Compound structural descriptors

From their secondary structure, we generated a vectorial representation of compounds using the molecular fragments obtained with ISIDA Fragmentor [16]. We generated all linear fragments of 1-7 atoms and all atom centered fragments of 2-4 atoms (ISIDA Fragmentor parameters -t3 -l2 -u7 -t6 -u4 -t0). We then coded each chemical compound as a binary vector, where each vector component represents the presence or absence of a given fragment. The vector length depends on the database analyzed (Supplementary Table 1).

In some cases, we found compounds with the same vectorial representation. These cases were merged, so that the dataset is non-redundant at the level of the vectorial representations.

### Scoring

From the vectorial representation of the compounds in a given pathway, we obtain a vector representation for that pathway (Pv) (Figure 1). Each vector component corresponds to a fragment in the database, and represents the probability (p-value) of observing it by chance in the compounds of the pathway, compared to a background distribution (all compounds in the database). These p-values are calculated with the cumulative hypergeometric distribution.

**Figure 1.**
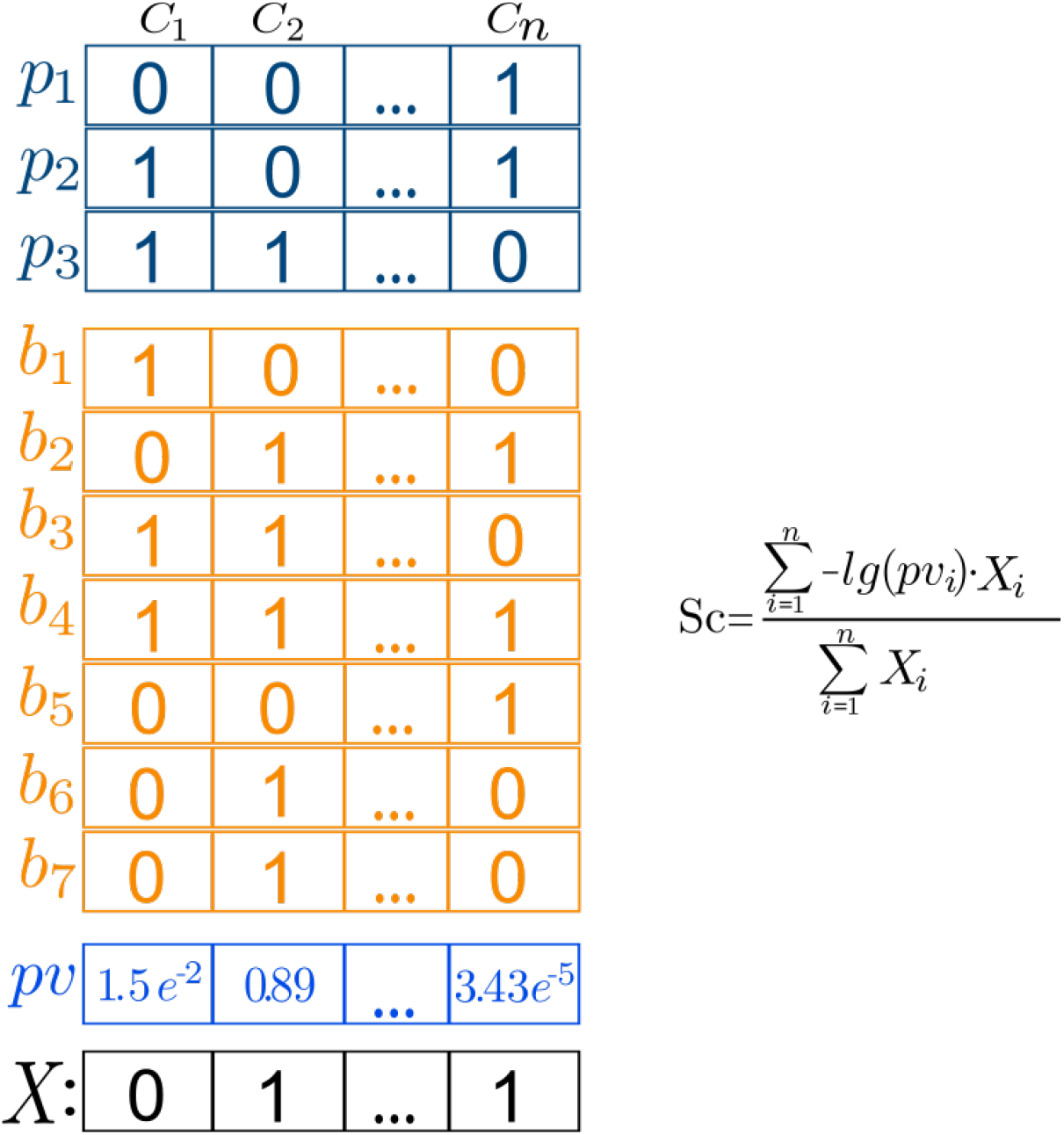
Example of calculating the score of a query compound (*X*) against a profile using iFragMent. Columns (*f*_i_) represent fragments, rows represent compounds: in blue (*p_i_*) those involved in pathway (P) and orange (*b_i_*) those that are not (background). Presence/absence is coded with 1/0. Vector *pv* contains the probability of observing a fragment in the compounds of the pathway by chance, considering their distribution in the background. The score of X against *P* (Sc) is calculated as shown.

With this pathway representation, we define a score for quantifying the matching of a given compound (*X*) against a pathway *P, Sc,* as:

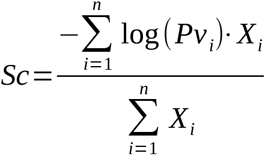

where *X_i_* is the ith component in the fingerprint representation of compound *X*, and P*v* is the probability vector of the presence of fragments in the pathway as described previously. To avoid log(0), those fragments with a p-value equal to zero were assigned a probability equal to the smallest p-value obtained divided by 50. Note that only fragments present in compound x (i.e. with a “1” in the vectorial representation) are taken into account with this formulation.

We observed, as expected, that scores Sc of compounds belonging to a pathway P tend to be higher than the scores of compounds not associated with that pathway (even if we exclude the compound to calculate Pv) (data not shown). However, scores of the same compound in different pathways are not comparable, so they can not be directly used as to predict in which pathway a compound participates. To solve this, we devised a random statistical model.

### Statistical model

In order to compare and rank the scores of a compound X in different pathways we calculate the corresponding z-scores using a null model, using a similar approach to [17]. For each pathway, we obtained a random distribution of scores (Sr). For that, we performed 100,000 randomizations of the matrix (NxM) of N compounds and M fingerprints (being N all the compounds in the corresponding database), and scored the resulting N randomized compound fingerprints against the pathway. Mean and standard deviations of Sr were obtained, and used to calculate z-scores from scores (Sc). We checked that Sr followed a extreme value distribution, as in [17]. Parameters characterizing each random distribution of scores (scale and location) were calculated with evfit function (MATLAB 2010b) and used to analytically calculate p-values from z-scores.

To predict the pathway of a compound, we calculate the z-scores and corresponding p-values of the compound against all pathway *pv* models. We then rank the pathway p-values by increasing value. We take those pathways in top-n ranking positions as the predictions, creating a whole family of predictors with increasing sensitivity and decreasing specificity. A ROC curve is calculated as n increases from 1 to the total number of pathways. Overall performance was quantified as the area under the ROC curve (AUC). A value of AUC = 0.5 would represent a random prediction (correct and incorrect pathways uniformly distributed in the ranked list), while values higher than that represent good predictions (correct pathways closer to the top of the list).

Both p-values and z-scores are reported by the iFragMent web server. The chemical substructures that are characteristic (enriched) in a biological pathway are obtained from the pathway Pv vector as those with the highest enrichment in that pathway (i.e. lowest p-value). These are used to highlight matched enriched fragments in the query compounds.

### K-nearest neighbour method

We implemented a k-nearest neighbour (k-NN) approach using the same vector representation of compounds. We calculate the structural similarity of a query compound to all the compounds in the database using the Tanimoto coefficient. We finally assign the pathways of the top-k most similar compounds. ROC curves and AUC values are calculated as previously explained.

### Evaluation

We evaluated our method, as well as the k-NN approach, with a 10-fold cross validation, using exactly the same partition of training datasets for both methods.

### Comparison with previous approaches

Dataset *human_unique* was downloaded from the Supplementary material of Marchiarulo *et al*. [8]. We used this dataset to perform a 10-fold cross validation of our method, and compare results with those reported in [8].

TrackSM software [10] was downloaded from https://dna.engr.uconn.edu/?page_id=648. TrackSM training dataset was obtained from the Config directory files, and were used to calculate iFragMent profiles, so that both methods can be compared trained in the same sets. Novel compounds in KEGG (release 83.0) not included in this TrackSM training dataset were used to predict pathways with both TraskSM and iFragMent.

## Results

### Overall performance

We tested the ability of our method to predict a compound’s biological pathway(s) as defined in four databases: KEGG pathway, Reactome, SMPDB and enviPath, which represent metabolic, signaling, and biodegradative routes among others. We use only those pathways with at least 10 distinct compounds.

To explore the largest number of chemical substructures, while at the same time not imposing any *a priori* chemical knowledge, we represented compounds as binary fragment vectors (see Methods for details).

We propose a method based on the probabilities of observing the presence of structural fragments in the compounds of a pathway just by chance. A compound having fragments enriched in a given pathway is a good candidate to participate in that pathway.

We evaluated the performance of the method using a 10-fold cross-validation approach (see Table 1 for overall performance results). Our method is a multi-label and multiclass approach, as it allows to assign several pathways to a compound (by taking the top-n predictions, see Methods). This allowed measuring performance by building ROC curves and calculate their AUC value (area under the curve), as we take as predictions from the top-1 to the full set of predictable pathways.

**Table 1.** Overall performance (AUC) of the method.

| Database | AUC |
| --- | --- |
| KEGG | 0.88 |
| enviPath | 0.90 |
| SMPDB | 0.74 |
| Reactome | 0.74 |

For all datasets we obtained good results (i.e. AUC >> 0.50). We obtained better overall results for enviPath (65 biodegradation pathways, 0.90 AUC), and KEGG (214 pathways, 0.88 AUC) than for SMPDB (333 pathways, 0.74 AUC) and Reactome (249 pathways, 0.74 AUC) databases. Neither the number of compounds nor the number of pathways was found to be related to these global performances. More details of each database can be found in Supplementary Table 1.

### Individual pathway performance

We found that results of individual pathways varied from random to excellent performance in all databases except enviPath (with AUC > 0.7 for all pathways) (Figure 2a).

**Figure 2.**
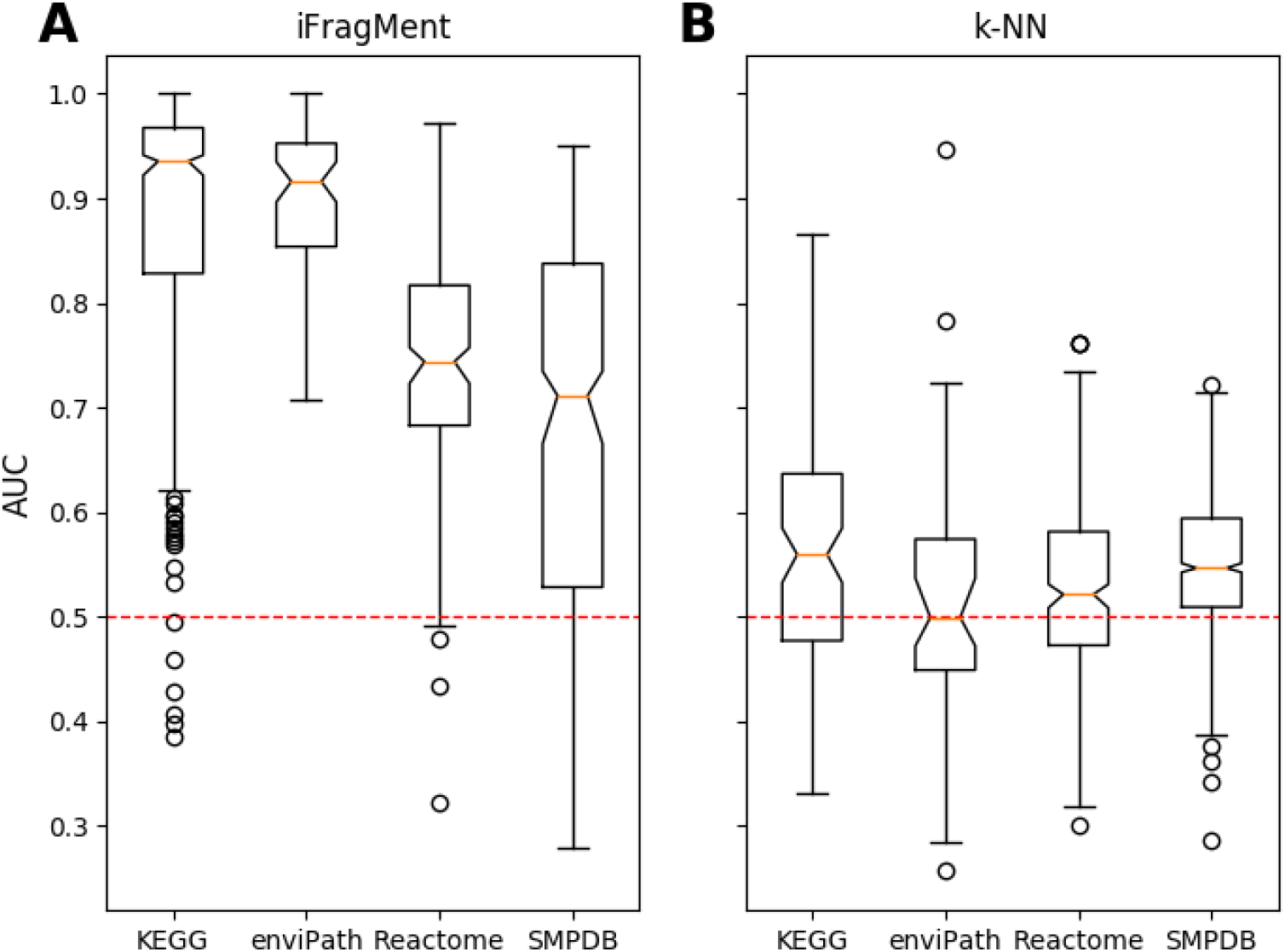
Distribution of AUC values for the individual pathways in the four databases tested for (a) iFragMent method and (b) k-NN method.

To compare our profile-inspired approach with a pair-wise one, we implemented a k-nearest neighbour classifier (k-NN) using the same vectorial (fingerprint) representation of compounds (see Methods). We calculate structural similarity with the Tanimoto coefficient, widely used in chemoinformatic approaches for structural comparisons of fingerprint representations. Our method attained higher AUCs in a 10-fold cross validation test than k-NN in the four databases tested (Figure 2). Details about each individual pathway performance using both methods are provided in Supplementary Tables 2.

In the following sections, we will analyse several factors that can affect prediction performance: pathway class, compound class (single or multi-pathway) and compound size.

#### Pathway class

We evaluated the performance of our approach for all KEGG pathways of a given general class (same top-level of the BRITE taxonomy) (Table 2). For that with constructed a ROC curve with the prediction results of all compounds associated with pathways belonging to that class, and calculated the AUC value. Some classes contained a limited number of pathways. Higher AUCs were obtained for ‘Drug development’ (1.00) (1 pathway: Histamine H2/H3 receptor agonists/antagonists), ‘Genetic Information Processing’ (0.94) (2 pathways) and ‘Metabolism’ (0.91) (153 pathways).

**Table 2.** Global performance (AUC for IFragMent and k-NN) for specific pathways within general KEGG pathway classes.

| Pathway Class | IFragMent<br>AUC | k-NN<br>AUC | Num.<br>pathways |
| --- | --- | --- | --- |
| Drug Development | 1.00 | 0.45 | 1 |
| Genetic Information Processing | 0.94 | 0.55 | 2 |
| Metabolism | 0.91 | 0.51 | 153 |
| Human Diseases | 0.82 | 0.52 | 10 |
| Environmental Information<br>Processing | 0.74 | 0.52 | 12 |
| Organismal Systems | 0.70 | 0.50 | 32 |
| Cellular Processes | 0.69 | 0.47 | 4 |

#### Compound size

We also observed that IFragMent performance varies largely depending on the number of chemical fragments present in a compound (a proxy of compound size) (Figure 3). AUC values of compounds with a small number of fragments (<32 for KEGG pathways), are very low in comparison with that of larger compounds. Similar results are obtained for the other three databases. Performance also decreases (although to a lesser extend) for very large compounds.

**Figure 3.**
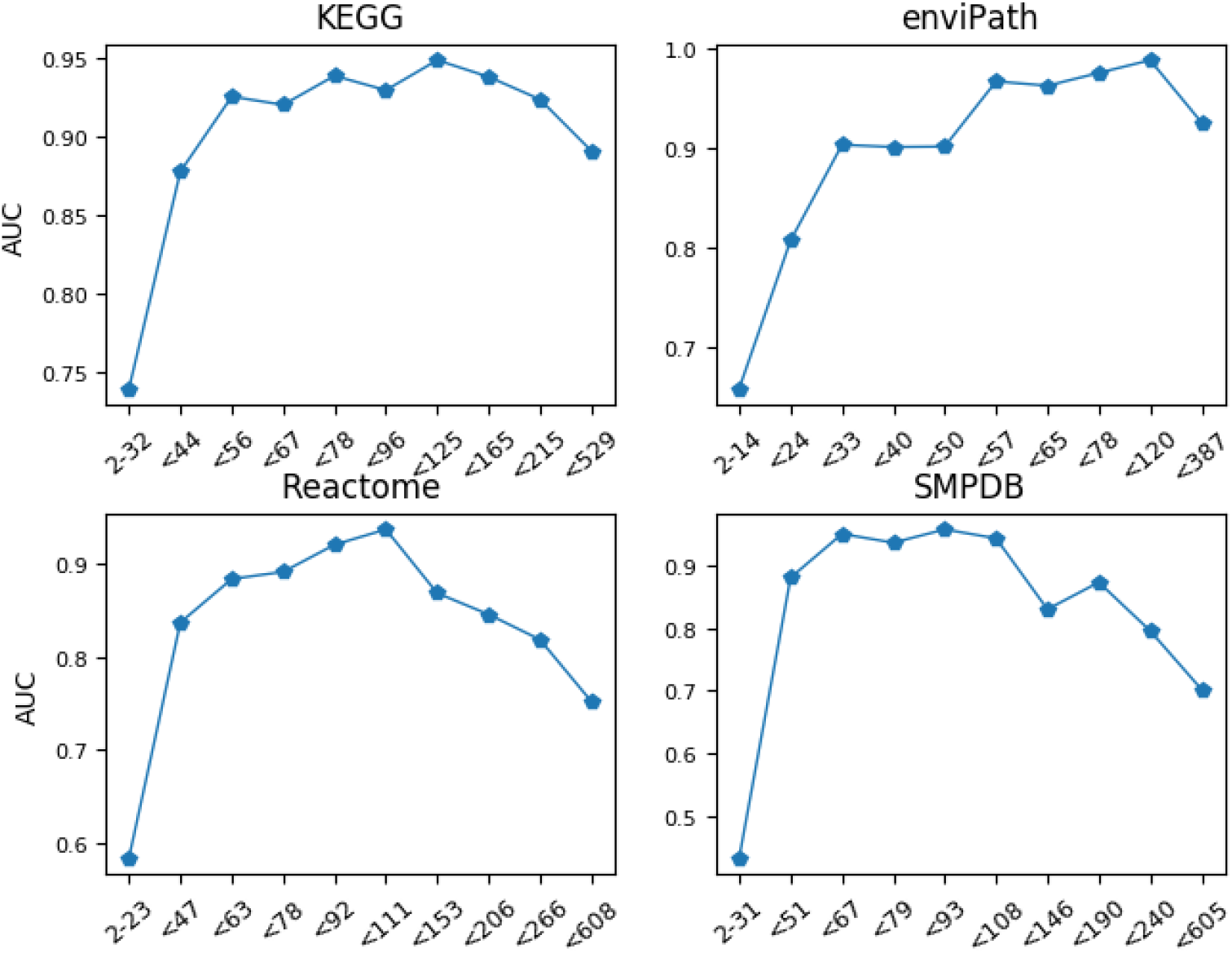
Performance of iFragMent (AUC) by number of fragments in the four databases tested. Intervals are defined to include a similar number of compounds.

#### Compounds in single vs. multiple pathways

We finally assessed the performance of compounds involved in a single pathway and those involved in multiple pathways. Compounds involved in multiple pathways achieved worse results than those involved in a single pathway (Table 3). E.g. for KEGG pathways, single-pathway compounds where predicted with an AUC of 0.95, in contrast to 0.83 AUC for multi-pathway compounds. As the fraction of multi-pathway compounds in each database varies, this partially explained the differences in global AUCs in the four databases studied (Table 3 and Supplementary figure 4).

**Table 3.** IFragMent performance (AUC) for compounds involved in a single pathway and those in multiple pathways

| Database | AUC uni | AUC multi | Single pathway compounds | Multi pathway compounds | %multi path compounds | AUC global |
| --- | --- | --- | --- | --- | --- | --- |
| KEGG | 0.95 | 0.82 | 4066 | 1170 | 23.2 | 0.88 |
| enviPath | 0.94 | 0.80 | 1254 | 91 | 12.1 | 0.90 |
| Reactome | 0.89 | 0.70 | 802 | 524 | 43.5 | 0.74 |
| SMPDB | 0.81 | 0.72 | 1062 | 346 | 35.9 | 0.74 |

### Comparison with previous methods

#### RF-Labute

To compare our approach with that of Machiarulo *et al* [8], which uses a Random Forest classifier and Labute descriptors for compounds (RF-Labute), we used their *human_unique* dataset (comprising 52 human pathways) and perform a 10-fold cross validaton of our method.

Neither overall performance measures nor numeric results were reported by Machiarulo *et al.* for their prediction of individual pathways (they did it only for the prediction of pathway classes). Instead, classification errors for individual pathways were graphically shown in a blox-plot organized in seven pathway classes (see figure 6D in original publication [8]). Assuming that this figure represents the out-of-bag estimate of error provided by RF, and that this error corresponds to the false discovery rate (FDR), we generated a similar figure for our results (Figure 4a). As RF-Labute is a multiclass but not a multilabel approach, we evaluated only the top-1 pathway prediction obtained by iFragMent for each compoud.

**Figure 4.**
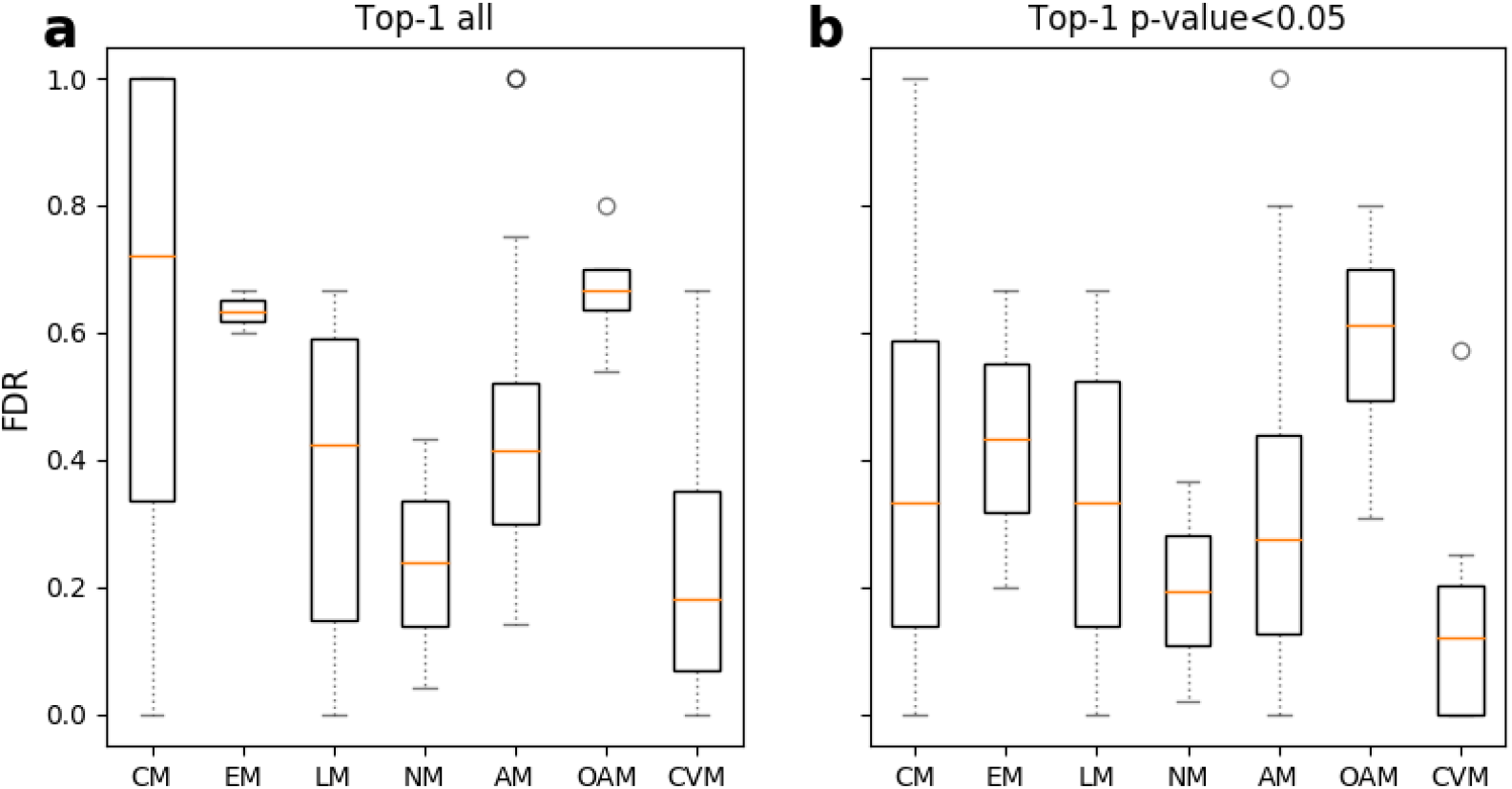
False discovery rate (FDR) of iFragMent using the *human_unique* dataset from Machiarulo *et al.* (a) All top-1 predictions. (b) Top-1 predictions with p-value <0.05

Visual comparison of both figures (Figure 2a and Figure 6D in [8]) reveals higher mean classification errors (FDR) of our method compared to RF-Labute for carbohydrate, lipid, nucleotide and amino acid metabolism pathways (CM, LM, NM and AM); comparable for both energy metabolism and cofactor and vitamins metabolism pathways (EM and CVM); and slightly lower for other amino acids metabolism pathways (OAM).

Part of the errors obtained by iFragMent in the *human_unique* dataset can be due to the small size of some of the pathways, for which reliable fragment statistics could not be obtained (15 out of 52 pathways contained less than 5 compounds, with more than 50% of them with less than 10 compounds).

In addition to the predicted pathway, iFragMent also provides an statistical estimate for each compound-pathway prediction (p-value). We have analysed the p-values obtained for each of the compound-pathway pairs (as defined in *human_unique*), and the rank at which they were predicted. As expected, classification errors (FDR) increase as p-value increases (Supplementary figure 1). Thus lower FDR (higher precision or PPV) can be obtained by p-value filtering, at the cost of decreasing sensitivy (TPR).

For example, considering all top-1 predictions iFragMent achieved a 0.66 TPR with a classification error (FDR) of 0.34. If we consider only the predictions with p-val <0.05 in the top-1 positions (Figure 4b), classification error (FDR) drops to 0.17, at the cost of lowering sensitivity (TPR) to 0.59. Thus, by filtering with p-values, we will miss some true compound-pathway associations (mainly those that are not based on enrichment of structural fragments), but make less mistakes.

In the previous section we compared iFragMent top-1 predictions with RF-Labute results. As our approach is a multiclass-multilabel classifier, it allows predicting more than one pathway for a compound. This feature can be exploited to increase the true-positive-rate (TPR, sensitivity). By considering the top-n predictions, TPR increases from 0.66 (top-1) to 0.79 (top-2) and 0.86 (top-3), at the cost of increasing also the number of false positives.

#### TrackSM

To compare our approach with TrackSM [10], we have designed a real world test scenario where new pathway-compound associations obtained from the release 83.0 of KEGG were predicted. To fairly compare methods, we use the same training dataset provided by TrackSM to construct the iFragMent chemical profiles. Hence, we trained TrackSM and iFragment with the same dataset, and generated predictions for compounds from a more recent version of KEGG. TrackSM reported errors for 41 compounds, that were not included in the comparison. We evaluated a total of 1313 compound-pathway associations involving 127 distinct pathway. Through the analysis of top-1 predictions we obtained a better performance with iFragMent (PPV = 0.41) as compared to TrackSM (PPV = 0.26). (See results in supplementary table 6).

### Chemical substructures associated to a biochemical pathway

Our method allows not only to assign compounds to their biological pathways (hence predicting their biological role) but also to detect the chemical substructures (fragments) that contribute most in such assignments (i.e. the matched components of the pathway Pv vector (Figure 1) that are statistically significant).

Figure 5 highlights the three fragments with lowest Pv values for the KEGG pathway “sphingolipid metabolism” (map00600) on the structures of the compounds associated to this pathway. It can be seen that these three fragments clearly delineate the the hydrocarbon chain and the polar head of sphingosine (C00319), while they are also highlighted in other compounds of the pathway.

**Figure 5.**
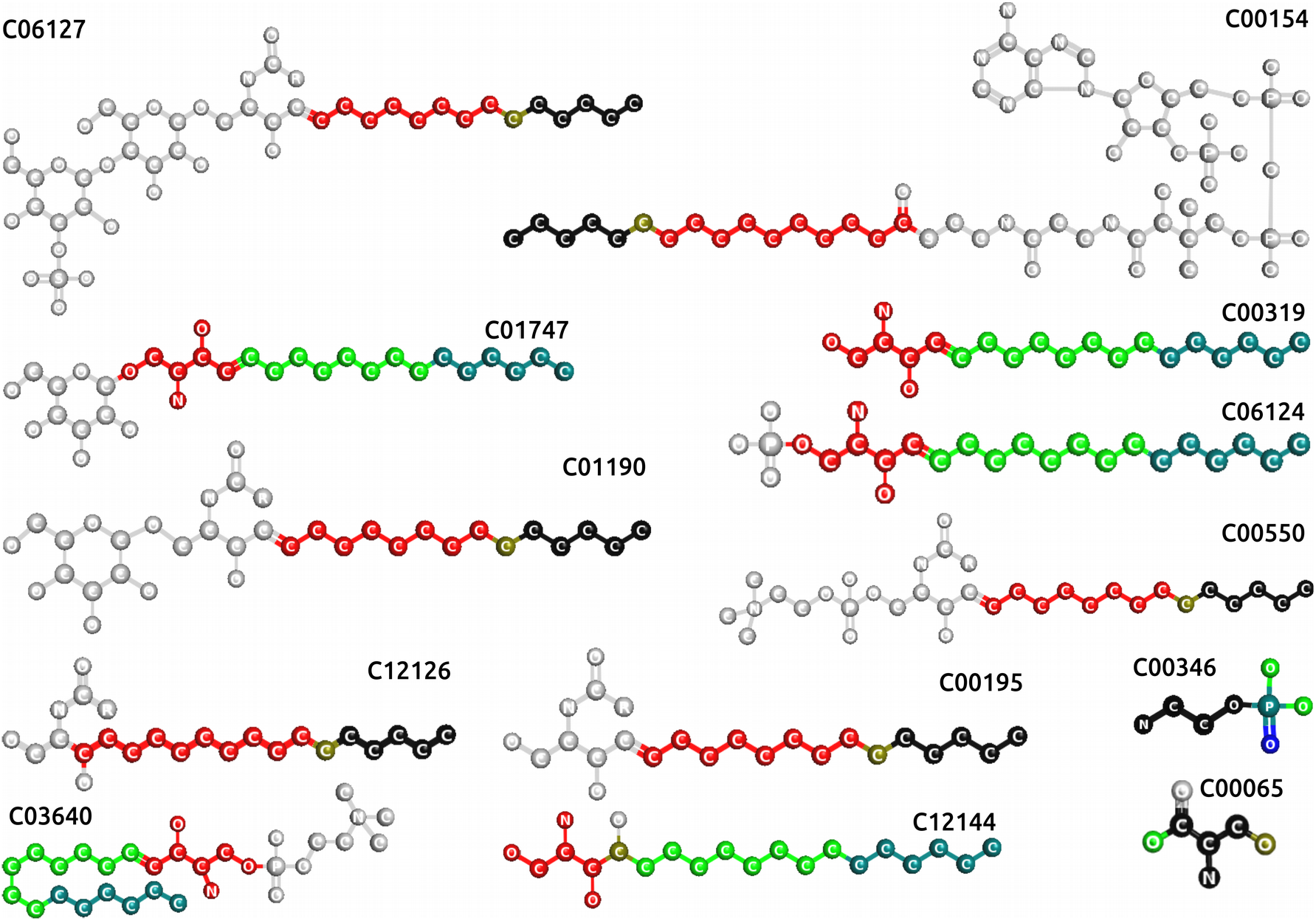
Enriched fragments (three lowest *pv* values) for the “sphingolipid metabolism” compounds (KEGG map00600). From lowest to highest, colored in red, green and blue; their intersection in teal (blue and green), olive (green and blue) and black (red, green and blue).

## Discussion

Profile methods are extensively used in the analysis of biological sequences (proteins, RNA and DNA) [18–21]. They are often used to infer the structure and function of an uncharacterized sequence by its similarity to a group of sequences (used to build the profile), as they detect remote homologies with greater accuracy than pair-wise similarities [22]. Sequence profiles are linear models, built upon the evolutionary relationships established among a group of biological sequences. Although we cannot establish evolutionary relations among chemical structures, chemical profile-inspired approaches have been successfully used in the classification of drug targets [23] relating receptors by ligand similarity, and the prediction of drug ‘off-targets’ [24] as compounds that bind to a protein have typically similar structures or substructures.

The ability of profile-inspired approaches to assign compounds to their biological pathways has not been explored. As metabolic reactions proceed step-wise, substrate-reactant pairs are structurally related. But given the linear or branched topology of reactions in pathways, we don’t know to what extend all compounds of a pathway are structurally related. Indeed, in some cases, structurally similar compounds tend to participate in the same metabolic pathway [25]. Additionally, the recent hypothesis of the “conquest of the chemical space” as the evolutionary driving force of the biological species [26] might suggest that evolution can also be indirectly reflected in chemical networks.

We demonstrate that a profile-inspired approach can be used to predict the potential biological pathway of a chemical compound from its 2D structure alone. We propose a method that relies on enrichment of structural fragments. Although performance varied largely depending on the pathway, we obtained good performances, especially for KEGG metabolic pathways and enviPath biotransformation pathways.

In this work pathways have been defined according to four public databases. KEGG, Reactome and SMPDB include not only metabolic but also other types of pathways such as disease, drug actions, transporting, signaling, etc. Yet, we found some general trends. In all databases, results for compounds involved in a single pathway are much better than for multi-pathway compounds. The percentage of multi-pathway compounds varied largely in the four databases, partially explaining differences in overall database performance. This could be due to the fact that multi-pathway compounds have “mixed” characteristics from more than one pathway, what might confound the predictor. This is equivalent to multi-domain proteins, for example.

Pathways for small compounds (e.g. < 32 fragments in KEGG) were identified with less accuracy than for larger compounds. This trend was observed in the four databases analyzed. This could be related to the absence of enough information in their chemical structures to discriminate the pathway they belong to.

The performance of our profile-inspired approach is higher than that obtained for a pairwise similarity approach (k-NN method) in all databases tested. We also compared our method with the two previously described approaches that addressed the prediction of individual pathways [8,10]. The three approaches differ both in the structural descriptors used to represent compounds and the algorithms. Our method (iFragMent) and TrackSM are multi-class and multi-label approaches, thus both enable the prediction of more than one pathway for a compound. Labute-RF can only predict one pathway for a compound (i.e. cannot handle multi-pathway compounds).

Testing a reduced set of single-pathway compounds involved in 52 human pathways, Marchiarulo *et al.* [8] reported lower classification errors than that obtained with iFragMent in its top-1 predictions. Half of the pathways contained less than 10 compounds (bellow the limit set in our work to obtain reliable statistical estimates of fragments). In contrast to machine learning approaches, like Random Forest, we establish a model beforehand to base our predictions: matching pathway-enriched fragments. This allows to provide an statistical estimate (p-value) to each prediction, which can be used to decrease classification errors (at the cost of missing some true positives).

In a real-world scenario, where newer compounds never seen by the systems were tested, iFragMent achieved higher PPV than TrakSM [10]. Our method does not consider pathway classes, while TrackSM predicts an individual pathway among those in a class previously predicted. Errors in pathway class prediction might limit TrackSM performance.

As we accumulate more data on the landscape of small chemicals underlying biological systems, new methodologies are required to mine it so as to extract useful information. Sequence profiles of biological polymers (DNA, RNA and proteins) are behind most approaches that allow mining and interpreting the massive genomic datasets. Consequently, we need similar approaches for the metabolism. A profile-inspired approach grouping functionally related metabolites is useful for assigning new metabolites to the group, as well as for detecting the structural fragments associated to that group (equivalent, for example, to the conserved/functional positions in protein profiles). The last allows getting insight into the chemical basis of the biological activity of a given group of functionally related compounds, in case it is unknown. Both, the assignment of chemical compounds to biological pathways and the detection of the informative fragments using this methodology can be performed by any interested user via the interactive graphical web interface developed.

## Acknowledgements

This work was partially funded by the Spanish Ministry of Economy, Industry and Competitiveness (MINECO) (project BIO2015-72091-EXP). J.L.I was the recipient of a pre-doctoral contract from the MINECO and European Social Fund (BIO2010-22109).

